# Neurocognitive Aging and Brain Signal Complexity

**DOI:** 10.1101/259713

**Authors:** Anthony Randal McIntosh

## Abstract

Brain organization can be appreciated across multiple spatial and temporal scales, where each scale affects the other in the emergent functions that we appreciate as cognition. As a complex adaptive system, the interplay of these scales in the brain represents the information that ultimately supports what we think and do. The dynamics of these multiscale operations can be quantified with measures of complexity, which are sensitive to the balance between information that is coded in local cell populations and that captured in the network interactions between populations. This local vs. global balance has its foundation in the structural connectivity of the brain, which is then realized through the dynamics of cell populations and their ensuing interactions with other populations. Considering brain function and cognition in this way enables a different perspective on the changes in cognitive function in aging.

Our initial work examined changes in brain signal complexity from childhood to adulthood. Across two independent studies, we observed an overall increase in signal complexity with maturation, which also correlated with more stable and accurate cognitive performance. There was some suggestion that the maximal change occurs in medial posterior cortical areas, which have been considered “network hubs” of the brain. In extending to study to healthy aging, we observed a scale dependent change in brain complexity across three independent studies. Healthy aging brings a shift in local/global balance, where more information is coded in local dynamics and less in global interactions. This balance is associated with better cognitive performance, and interestingly in a more active lifestyle. It also seems that the lack of this shift in local/global balance is predictive of worse cognitive performance and potentially predictive of additional decline indicative of dementia.

---

The main goal of my chapter is to convince you that age-related changes in brain function can be best appreciated from the perspective of complex network dynamics. The brain is a collection of networks that are constantly changing their interactions. The dynamics have spatial and temporal signatures, which are vital to the emergence of stable behavior. By this same token, brain dysfunctions can affect space and time, which can provide new avenues to consider in diagnosis and treatment in the face of disease or damage.

Our brain is a complex adaptive system, showing constant flow of activity and interactivity during coordination of behavior and cognitive function. Complex adaptive systems can show an optimal balance of integrating and segregating information (Tononi, Sporns, & Edelman, 1994). In the brain, high complexity arises from the interplay of brain structure and function (Sporns, Tononi, & Edelman, 2000a, 2000b). Changes in this interplay underlie cognitive evolution across the lifespan (McIntosh et al., 2010), while brain damage or disease disrupts the interplay leading to cognitive dysfunction (Fornito, Zalesky, & Breakspear, 2015).

One feature that contributes to brain complexity is the space-time structure of anatomical connectivity (Figure 1). From a general anatomical perspective, neighboring elements are more densely connected relative to more distal elements. For the local connections, impulse transmission is very rapid and effectively instantaneous, while impulses from distal regions will arrive with differing delays based on the relative distance and conduction velocity. In the absence of dynamics, the space-time structure conveys the *potential* network configurations that can be expressed (Deco, Jirsa, & McIntosh, 2011). The space-time structure matrix, when extended into three dimensions, provides a useful way to visualize which connections could be active at a given a point in time.

**Figure 1.**
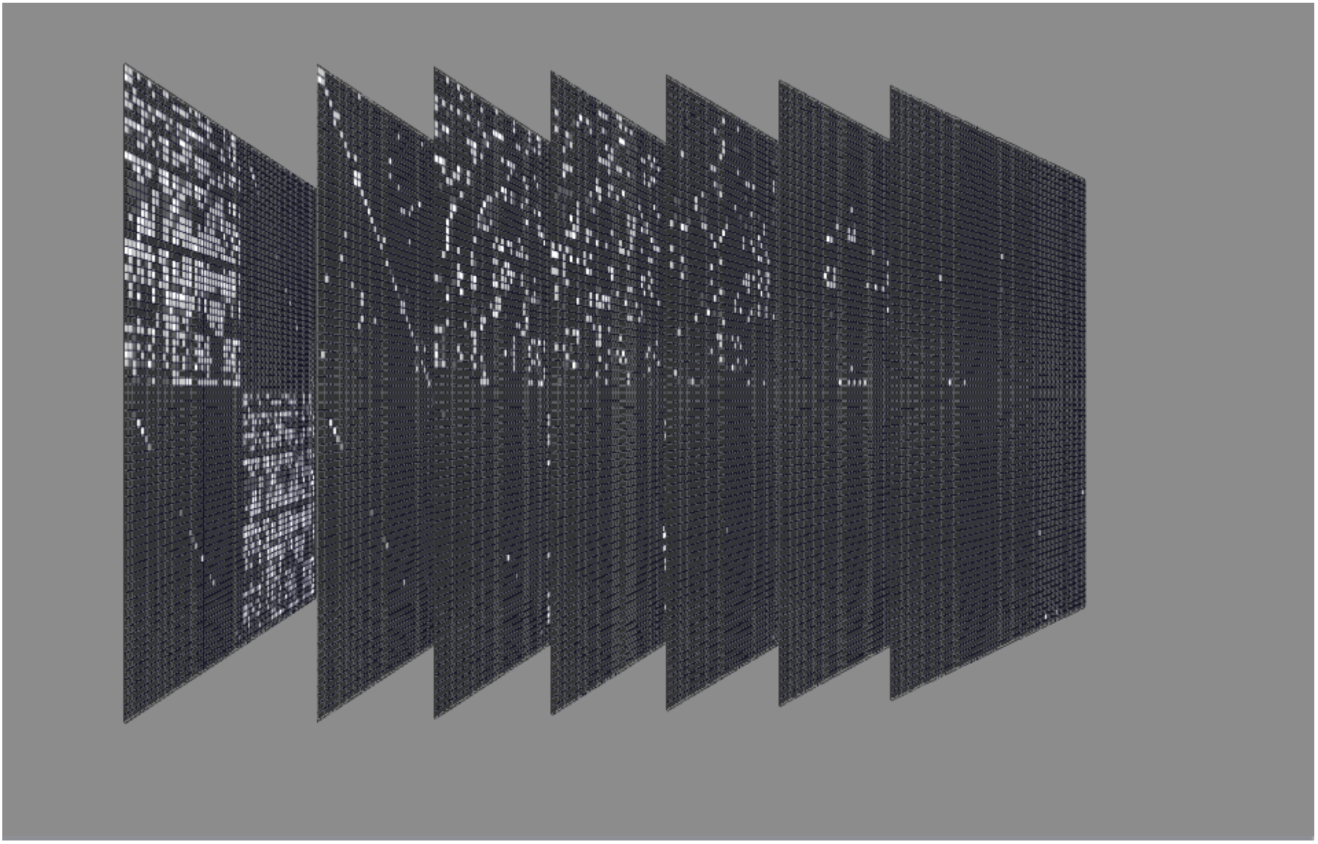
Space-time structure of anatomical connectivity. The first matrix on the left is the adjacency matrix showing the connections among 76 bilateral region of interest, where each cell in the matrix is a region. The top left quadrant is the left hemisphere and bottom right the right hemisphere. The other two quadrants represent cross-hemispheric connections. Shaded cells indicate the presence of a connection, with the intensity indicating connection strength. Subsequent slices show the adjacency matrix reconfigured to illustrate the time it takes for an impulse to traverse a connection based on strength and conduction velocity. Slices indicate connections that could be “active” at a given time step, from 0 ms to 150 ms in steps of 25 ms.

The space-time structure conveys the potential network configurations, but it is the dynamics that actualizes them. The dynamics in complex systems, like the brain, show varying degrees of ***noise*** that is, paradoxically, vital for its function (Faisal, Selen, & Wolpert, 2008; Kosko, 2006). Noise in the brain manifests at two levels: local cellular operations and network dynamics. The local operations show a level of stochasticity such that there are imprecisions in biophysical operations (e.g., channel openings, ion exchange). The level of local noise is seemingly “tuned” so that signal propagation can be maintained and even facilitated. A good example of this is the classic stochastic resonance phenomenon wherein nonlinear systems with an optimal amount of noise are able to detect weak signals (Kosko & Mitaim, 2003; Manjarrez, Rojas-Piloni, Mendez, & Flores, 2003; Ward, Neiman, & Moss, 2002; Wiesenfeld & Moss, 1995).

The local noise affects how incoming signals are received. The local noise, on top the time delays in the space-time structure, combine to what can be considered ***network noise*** leading to variation of functional network configurations. As the number of functional networks increases with maturation and experience, the network noise reflects both the number of potential configurations available and those that are instantiated (Ghosh, Rho, McIntosh, Kotter, & Jirsa, 2008). In the case of a completely deterministic system, the network trajectories are fixed and hence no matter what the signal is, the information conveyed by the network fluctuations is the same. In the face of a space-time structure with variable time delays and local noise, the information capacity will be higher reflecting the multiple network configurations that can be realized. Thus, while noise has a negative connotation, in a nonlinear system, noise serves a crucial role in enabling the temporal evolution of the system. This property was acknowledged at least 50 yrs. ago (Pinneo, 1966), and has seen a revival as part of the explanation for the ubiquitous “resting-state networks” that have been characterized in a plethora of functional neuroimaging studies.(Damoiseaux et al., 2006; Deco et al., 2011; Fox et al., 2005).

In the brain, signals from the exact same network may show different propagation patterns based on the history of the system (local dynamics), which can redirect functional network configurations. This is a hallmark feature of complex adaptive systems that constantly explore current configurations, but also assess new ones in case they result in better solutions for the system.

The evolution of network dynamics is believed to relate to the similar evolution of cognitive and behavioral functions. From this link, it would be reasoned that as the richness of network dynamics grows, so too does the richness of the cognitive functions. The relation would not be completely linear, however, where at some point excessive network dynamics would be unable to support a coherent cognitive flow. This classic “inverted-U” relationship also illustrates the balance that must be maintained between segregated and integration operations (or local vs. global). To echo a previous point, the brain moves between integrated and segregated states to optimize the integration of information, or complexity.

## Measuring Brain Complexity

The quantitative estimation of complexity in the brain has been an active area of study for at least 20 yrs. Entropy, as defined by Shannon (Shannon, 1948), is a useful metric as it gauges the predictability of a signal and the information content. For example, a sinusoid carries little information once the frequency and amplitude are known, but a multi-frequency signal requires more measures to completely characterize it. Hence, a sinusoid would less entropy than a multi-frequency signal.

Tononi and colleagues (Tononi et al., 1994) derived a measure of brain complexity from entropy, reformulated to reflect the balance of integration and segregation of brain dynamics. In this instantiation, systems with high complexity would show maximal information integration. The formulation emphasizes the interactions between networks and specifically the shared information between elements, known as *mutual information.* The challenge with this formulation is the practical application to empirical data, as the estimation of complexity requires an exhaustive assessment of network partitions and the mutual information between such partitions.

An alternative is to consider approximate methods that are sensitive to the shared information between neural elements. Linear estimators are good for this, such as correlations of coherence estimates, but are limited in the network dynamics that can be measured. In particular, higher-order nonlinearities may be critical for transition of network configurations, but would be invisible to linear estimators.

Entropy alone could be a useful measure, but a pure noise signal can also show high entropy (Costa, Goldberger, & Peng, 2002a). In this case, its important to measure whether the information conveyed is the same across scales, in this case time scales. The scale-dependency is a critical factor in such a differentiation, which links back to the space-time structure mentioned above. One metric that accounts for timescale dependency is multi-scale entropy (MSE (Costa, Goldberger, & Peng, 2002b, 2005)), where entropy is estimated across progressively coarser time scales. (Entropy is estimated using Sample Entropy (*Se*, (Richman & Moorman, 2000)), which operates well on discrete timeseries that are common in empirical data. The expectation is that signals with little information will show a rapid decline in entropy with timescale reduction. So too, pure noise will show a decline in entropy with timescale. This is because pure noise has no timescale dependency. Truly complex signals (e.g., colored noise) have dependencies that can arise for the space-time structure, and thus will show high entropy across multiple timescales.

The remainder of this chapter will focus on how we have used the principles of complexity to assess the changes in information processing capacity in development and aging. The hypothesis is that if development represents an increase in the information processing capacity the brain, the MSE should increase with maturation. In aging, the simple hypothesis would be that information processing capacity goes down and thus so should MSE. However, in both development and aging, there would very likely be scale-dependency in how these changes manifest. In development, there are multiple local and distributed network changes that reflect both the biological maturation and the effects of experience. In aging, biological changes are more subtle, as are the cognitive changes, and hence the MSE changes may be less extensive than those observed in maturation. Finally, we will touch on clinical applications of MSE, which indicates that there may be prognostic utility in the scale-dependency information.

## Applications

## Maturation and Brain Complexity

Our first examination of the changes in MSE focused on development (McIntosh, Kovacevic, & Itier, 2008). Here we used EEG data recorded while kids 8–15 yrs. viewed individual faces in a 1-back memory task (i.e. “Does the current face match the one you just saw?”) Young adults from 18–25 yrs. were also tested in this paradigm. The evoked-potentials to the faces showed a well-characterized developmental change where the youngest child showed a high amplitude response initially, without much beyond that (Figure 2). With maturation, the overall amplitude of the response reduced and secondary responses became more evident. This change in the mean response signal is consistent with a move from a deterministic system, where there is a large stereotypic response on each trial, to a more stochastic system with more complexity reflecting multiple network dynamics.

**Figure 2.**
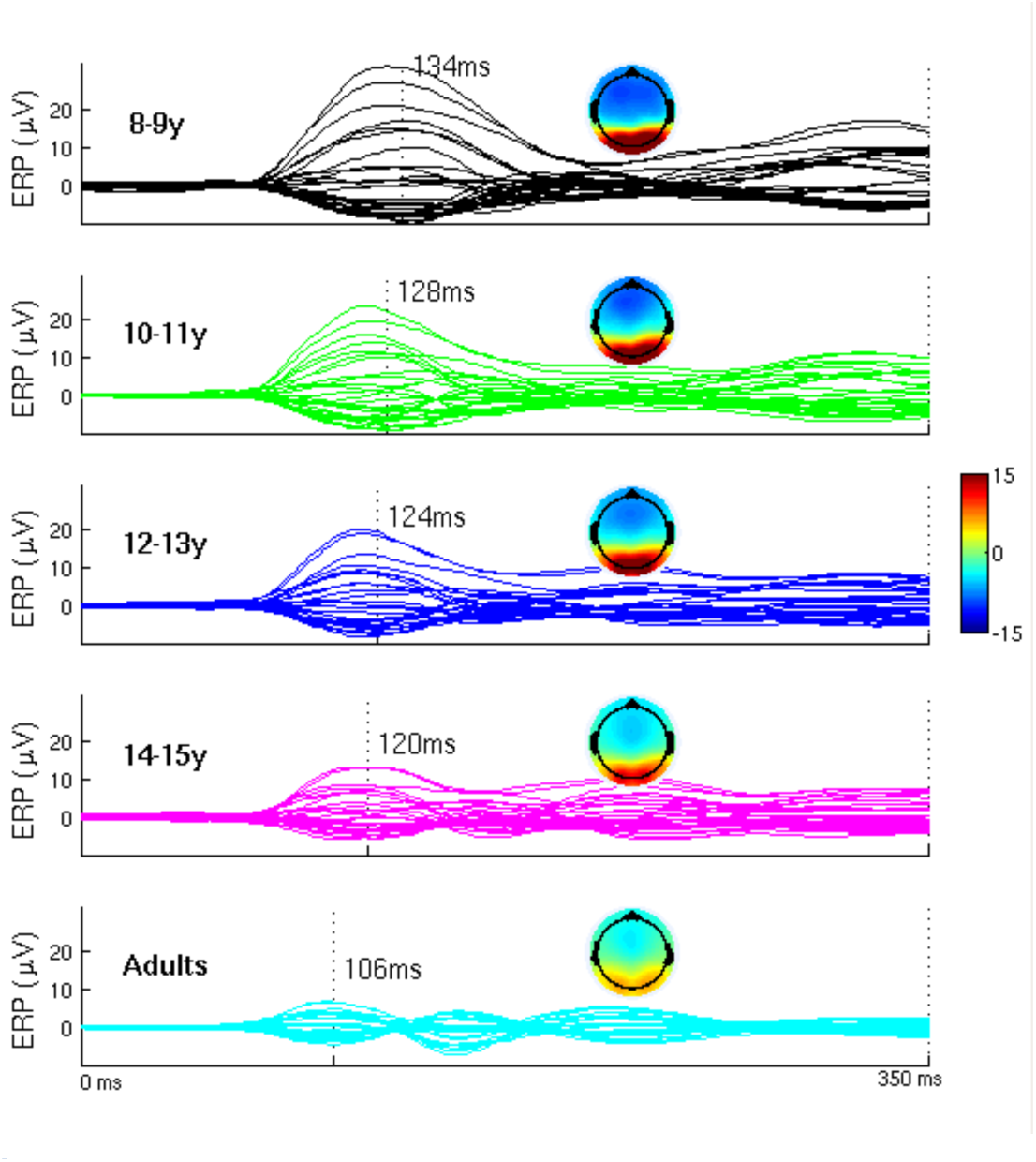
Average evoked-potential within age groups in response to a face. “Butterfly” plots show the time series for each electrode across the 350ms trial. The vertical mark indicated the approximate time of the classic P100 response, and is displayed on the topoplots for each group. The youngest group has the slowest peak response, but the largest amplitude. The shifts with age where by adulthood the peak is faster and the evoked response has a more multi-componential feature.

The increase in variability was reinforced in an assessment of single-trial variability within each subject using principal components analysis (PCA). The assumption underlying the PCA application was that if there is more variance in the trial-by-trial response, this should come out at greater dimensionality where more components would be needed to capture an equivalent proportion of variance across persons. Indeed it was the case that with maturation, there was an increase in the number of components needed to capture 90% of the variance in a subjects data (Figure 3). Another feature captured in the PCA was the dimensionality reduction from the stimulus onset. Pre- vs. post-stimulus PCA was compared and, while there was a reduction across all ages after stimulus onset, the magnitude of the change reduced with age. This would be expected if maturations moves the brain from a deterministic system to one with higher complexity, as a deterministic system would show greater overall entrainment from a stimulus.

**Figure 3.**
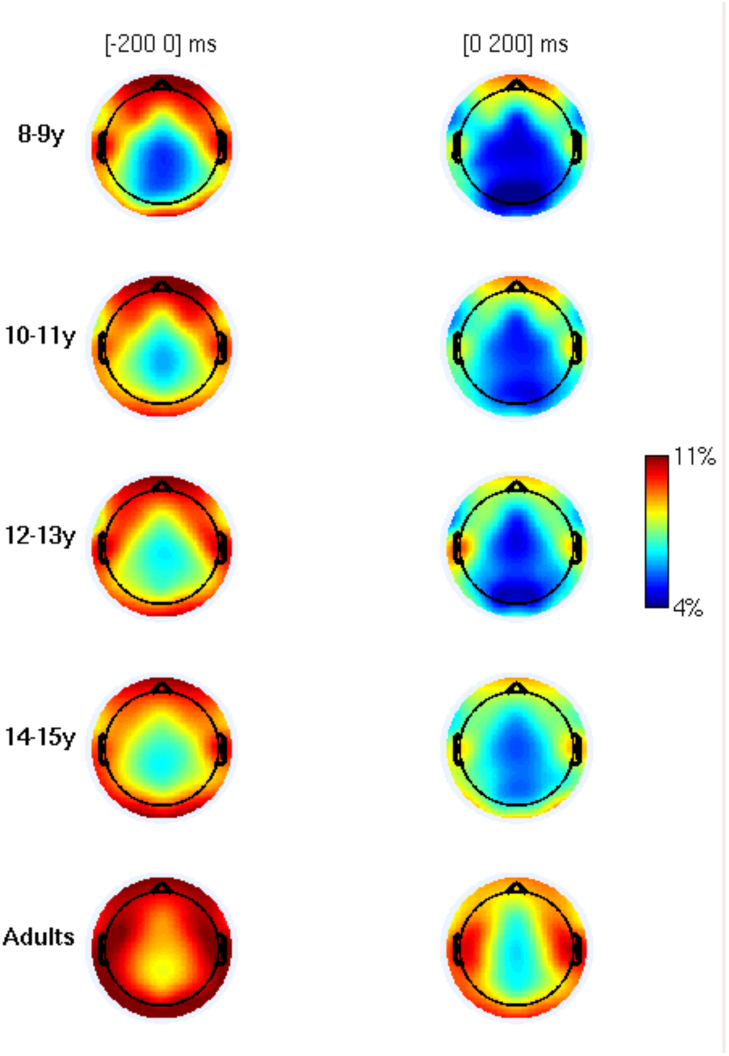
Topoplots for each EEG channel within age group showing the number of principal components (%total) need to capture 90% of the total variance across single-trials. The PCA was done both for pre- (-200 to 0) and post-stimulus (0-200ms) trials. In additional to higher dimensionality in general with increasing age, the dimensionality reduction from the stimulus also decreased with age.

As might be expected, analysis of MSE also showed a maturational increase. While the curves suggest a general increase across all scales, there is a somewhat stronger effect at the mid-range of the timescales (scale 6–10). The curve for adults shows the highest entropy scales and the children fall ordinally in place with the youngest kids at the bottom of the graph (Figure 4).

**Figure 4.**
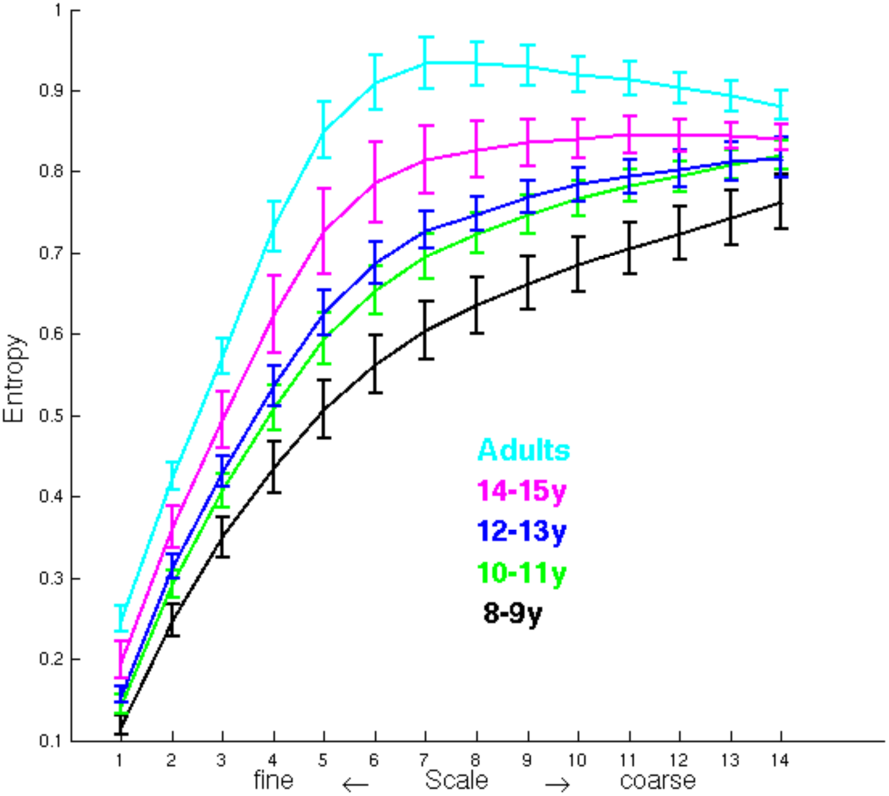
Mulitiscale Entropy (MSE) curves for each age group at a posterior EEG channel. Sample entropy (Entropy) is plotted from fine to coarse time scales. Maturation shows a general increase in MSE, with the largest change at mid-range scales (6–10).

While the observations of increasing signal variability and complexity with maturation are intriguing, relating these changes to behavior helps with interpretation. We correlated the PCA and MSE metrics with reaction time, the coefficient of variation of reaction time (SD/Mean, CVrt) and accuracy on the 1-back task. First, mean reaction time did not show a stable correlation across the entire sample. However, CVrt showed a negative correlation with PCA and MSE, indicating that a steadier reaction time was related to higher signal variability and complexity. Accuracy showed a positive correlation, where greater accuracy was correlated with higher signal variability and complexity. Said together, brains with greater noise show more stable and accurate behavior. It is noteworthy that this pattern remains, albeit somewhat weaker, if age is regressed out the sample, indicating that this relationship may be a general feature, rather than an exclusive reflection of development.

Two issues are noteworthy here regarding the metrics of brain noise. First, it is unlikely that these effects can be explained by **measurement noise**, particularly in the case that the MSE measure is able to differentiate structured noise from random/white noise. One may expect that measurement noise would be higher is children, which is opposite of what we observed with PCA and MSE estimation.

Second, spectral power analysis showed complementary effects to that of MSE, with the highest age-related changes in the beta bands (15-30 Hz). Thus, the temptation is to explain the MSE results as a simple reflection of spectral power changes. Insofar as spectral power and MSE will reflect similar underlying biophysical phenomena, this is a reasonable explanation. Local estimates of spectral power will reflect the influences of local (high frequency) and distributed (low frequency) sources. MSE, as a local measure, will also reflect local (fine scale) and distributed (coarse scale) influences. Both capture the capacity inherent in the space-time structure. Where the two measures diverge is how they reflect the interdependencies across space and time. Spectral power decomposition is not sensitive to the cross-spectral dependencies, whereas MSE estimates are. This difference is most easily captured in simple simulations (McIntosh et al., 2008). We can transform the adult spectral power data to match that of the children. MSE was calculated on the reconstructed data, and compared to the original curve. Here the MSE curves in the modified data showed a similar profile as in the children, with lower entropy at mid-scales. Another simulation was done to destroy the cross-spectral dependencies. Here the signals from the adults were decomposed into the Fourier domain, and rather than modifying the power of the coefficients, the phases of the frequencies were scrambled, which leaves the spectral power distribution the same, but changes the higher-order dependencies. As before, the signals were reconstructed and MSE calculated. The MSE estimates were altered showing *higher* entropy in the modified time series. These simple simulations indicate that spectral power and MSE provide complementary characterizations of brain signals.

### Replication

Replication is a crucial aspect of scientific investigation. We were able to replicate the developmental patterns in a second study, which used MEG rather than EEG in the same 1-back task (Misic, Mills, Taylor, & McIntosh, 2010). These data also covered a broader age-range: 6 to 16 yrs.

The MEG data were source-modeled, expressing the signals as emerging from cortical sources rather than from the scalp surface. MSE analysis of these data showed the same age effects as we noted earlier – an increase in MSE with age. The effect was present across most sources, but was strongest in medial parietal cortex. Interestingly, this region has been labeled as a network hub, in that its structural connectivity pattern provides capacity to interact with many different networks (Hagmann et al., 2008). Such a region could be expected to show higher complexity.

Brain-behavior analysis also replicated what we noted before, though this time reaction time was also correlated with MSE, where faster and more consistent reaction time was related to higher MSE. These relations did not show regional specificity.

Spectral power analysis of these data showed a similar overall effect to what we found previously, with the greatest changes in the middle beta band (15–20 Hz). What is interesting here is the distribution of the maximum spatial effect differs from MSE. Spectral power effects were most prominent in lateral parieto-occiptial cortices.

### Interim Summary

In development, the brain increases its overall capacity for information processing. There are a multitude of changes: synaptic proliferation and pruning, white matter connectivity increases, and various environmental effects that interact with the emerging cognitive function. Such a broad range of effects may be expected to increase brain noise across many scales, which are well-reflected in the MSE estimation. We also found that these general changes can be mapped as far back as 1-month-old infants (Lippe, Kovacevic, & McIntosh, 2009). Importantly, the MSE changes are correlated with behavior, suggesting a strong link to signal complexity and information processing capacity.

#### Aging

Complexity in the brain shows a gradual increase across most scales in development. In aging, one could hypothesize a few scenarios. Given the proven decline in overall cognitive function, one may expect complexity to go down. On the other hand, the cognitive changes are not as broad as one sees in development, in that some functions do indeed decline, while others remain relatively stable or perhaps even increase with age. Hence, it may be that the changes in brain complexity observed in normal aging show more spatiotemporal dependency than in maturation.

Our first exploration was a study that examined two datasets, one EEG and one MEG(McIntosh et al., 2014). The EEG data were acquired in three age groups, young (mean age = 22 yrs.), middle (mean = 45 yrs.) and old adults (mean = 66 yrs.), while performing visual perception tasks. The MEG data were collected from two groups (young: mean = 23 yrs.; old: mean = 70 yrs.), performing an auditory-visual attention task. The data were source modeled to allow more inferences on regional changes.

Both data sets showed essentially the same temporal effects, suggesting that the nature of age-related changes cannot be characterized as simple increases or decreases of signal, or signal complexity, but rather the effects depend on “when” and “where” you are looking.

The MSE changes in the EEG data nicely illustrate this point (Figure 5). What we saw in the group average curves is that the finest scales, the middle and old-aged group show higher Se, but the curves cross at middle scales such that at the most coarse scales these two groups show lower Se compared to the young group. There was regional variation in the strength of this effect, with strong precuneus and left prefrontal involvement (Figure 6). The MEG data showed the same temporal effect with the older group having higher Se at fine scales and lower Se at coarse scales. The effects were broad and showed less regionality than the EEG data.

**Figure 5.**
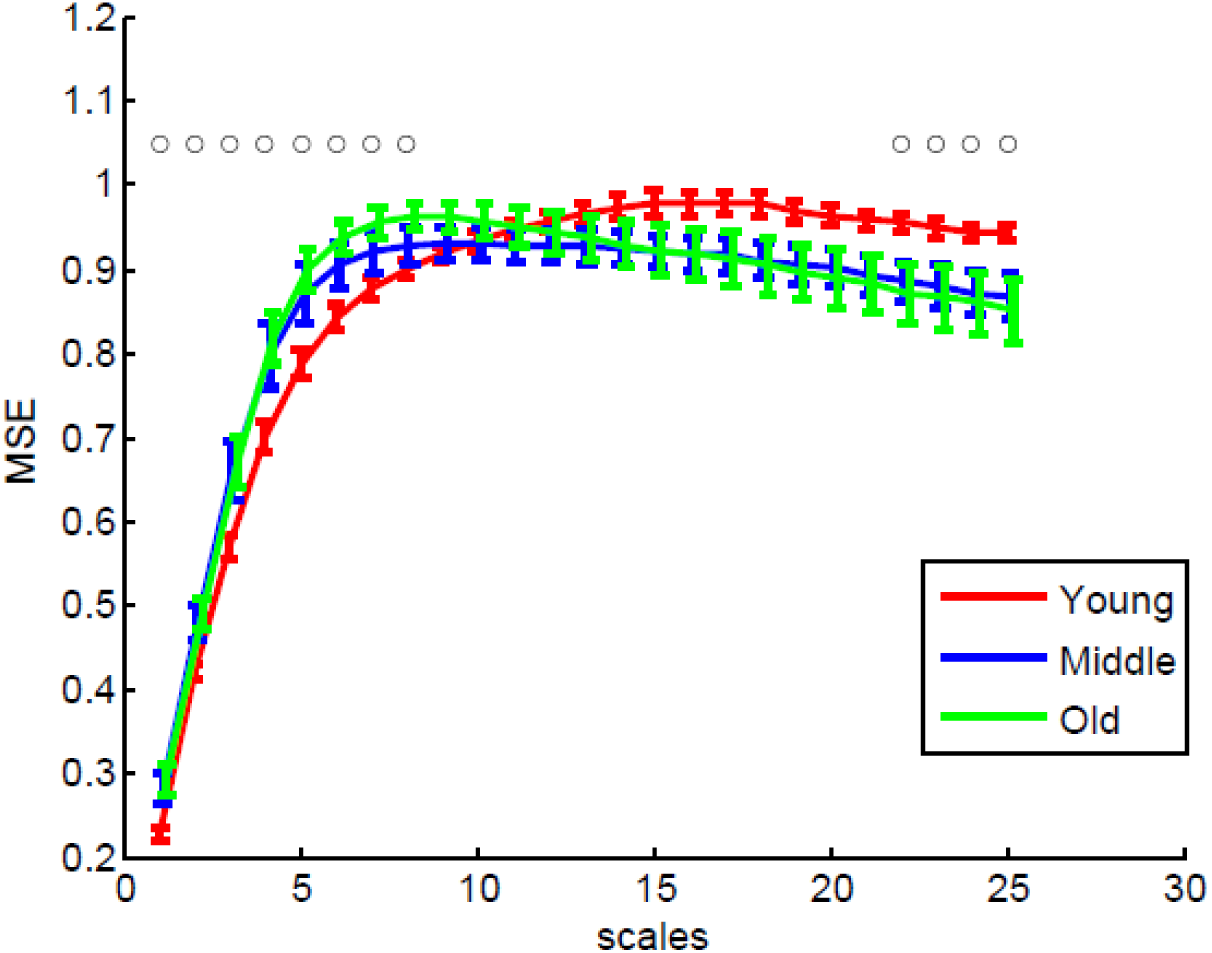
Mean (+/- SE) MSE curves for the three age groups at a precuneus EEG source. Open circles above scales indicate points of significant divergence of curves from one another, as assessed by multivariate partial least squares analysis.

**Figure 6.**
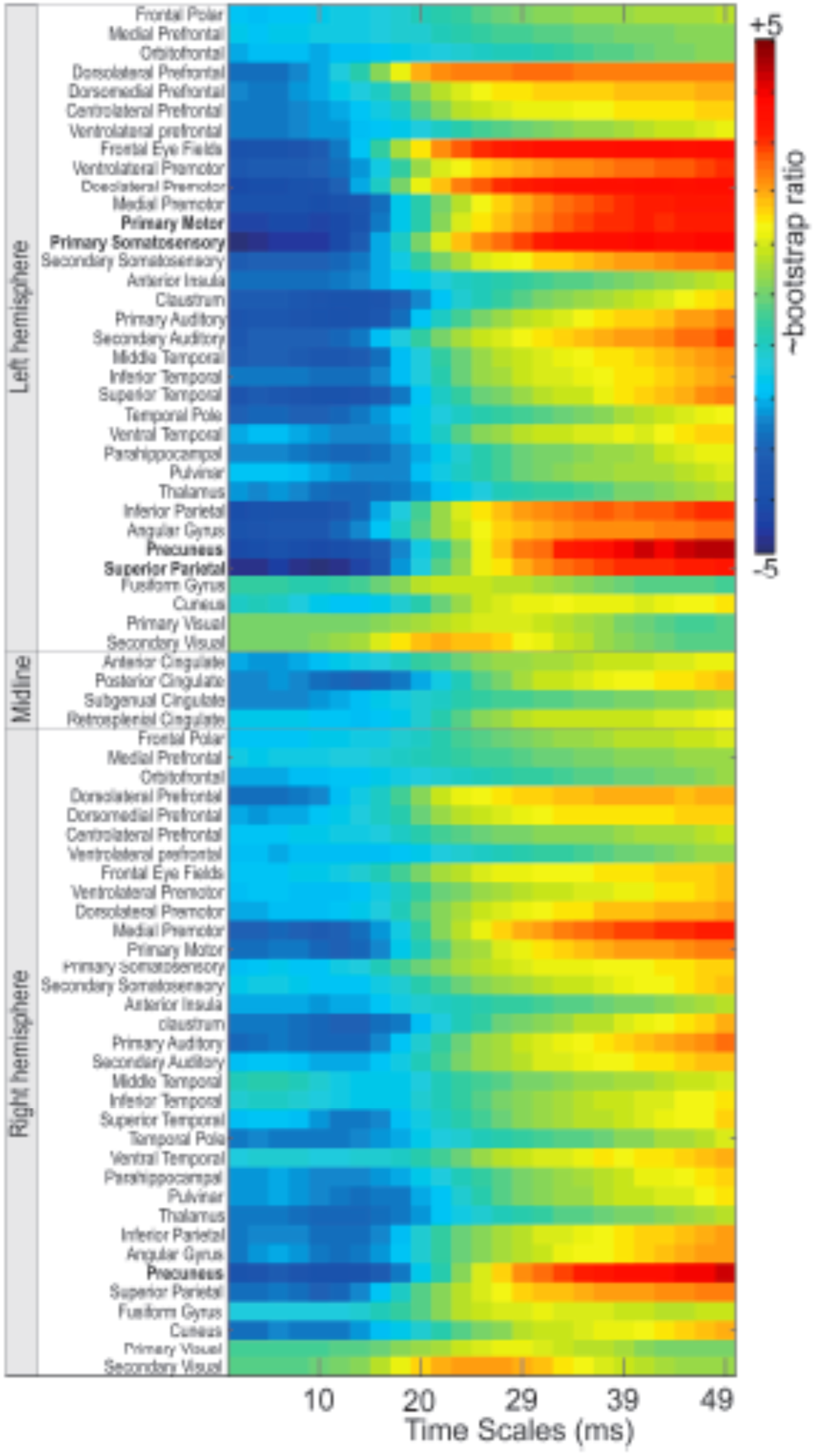
Summary of EEG sources and MSE time scales showing group differences (Figure 7). Blue colors indicate higher entropy in middle and older adults, while yellow and red indicate higher entropy in younger adults. Bootstrap ratios are comparable to z-scores and are derived y bootstrap estimation within the partial least squares analysis framework.

As noted earlier, MSE is effectively a local measure in that it captures the variation in Se from a given location without considering the source of the variation. Specifically, the activity in an area will be a function of local dynamics from the interacting cell populations, and the influence of other connected areas. The MSE estimation will reflect both of these effects. While there are more recent advances in MSE that allow multivariate calculations (Ahmed & Mandic, 2011), with these data we were able to parse the MSE effects into those related to local dynamics, and those related to distributed effects.

This parcellation can be captured by considering a Venn diagram for the different sources of entropy (Figure 7). The total entropy of a regions signal (H(x) and H(y)) is a function of local variation and interactions with other regions. The interactions can be estimated using mutual information (I(x;y)), which is the shared Se, when Se unique to each source is parsed out (H(x|y) and H(y|x)). Similarly one can estimate the local Se, which would be the Se for a given region (again H(x|y) and H(y|x)), with shared Se from all other sources parsed out. Like MSE, this estimation can be done for each scale.

**Figure 7.**
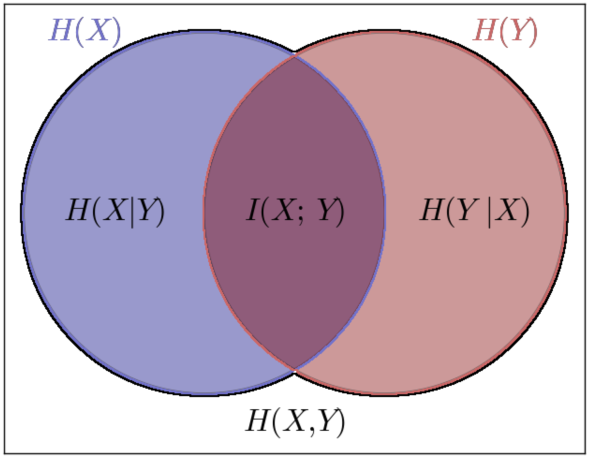
Partitioning of entropy (H), into local and global effect. H(X) and H(Y) are total entropy for variables X and Y, and H(X,Y) are the joint entropy for both. Mutual information (I(X;Y)) is the uniquely shared entropy between X and Y, while H(X|Y) and H(Y|X) are = entropy estimates for X and Y, conditioned by the entropy of the opposing variable. I(X;Y) is distributed entropy, and H(X|Y) and H(Y|X) are local entropy

When expressed in terms of local and distributed entropy, a clear picture emerged that helps explain the effects noted in the MSE estimation (Figure 8). Local entropy for both the MEG and EEG data set was higher for the older groups. Distributed entropy showed a predominant decrease for the older groups, with the largest effects involving cross-hemispheric interactions.

**Figure 8.**
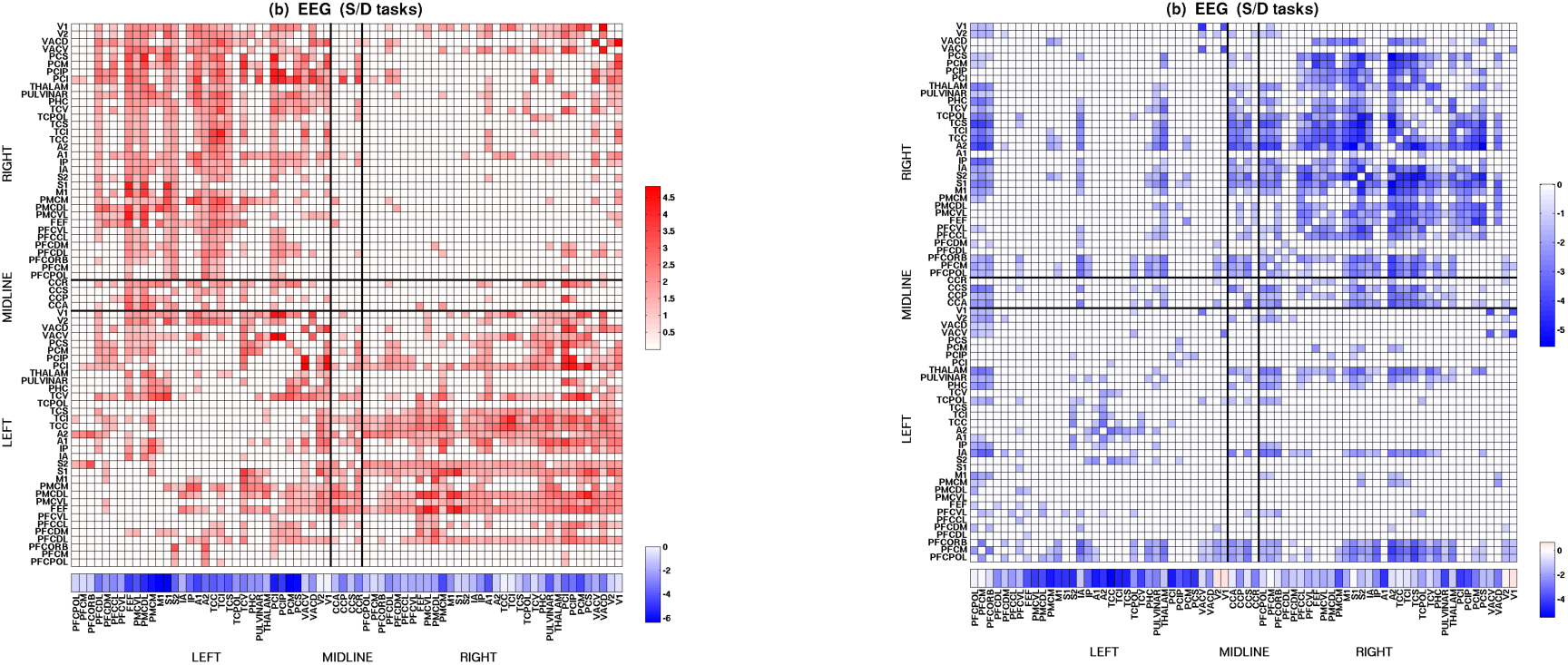
Matrix showing the distributed entropy decreases (red) and increases (blue) for the interactions between EEG sources in healthy aging. Upper left and lower right quadrants are cross-hemispheric interactions and upper right and lower left are within hemisphere. Aging shows strong cross-hemispheric decreases in distributed entropy and increases within hemisphere. The vector below the matrix indicates local entropy, which shows general increase in aging.

Some clarity to the picture now emerges with the analysis of local and distributed effects, considered with the MSE analyses. One can interpret the changes are reflecting age-related changes in network dynamics such that a larger proportion of information is being processed locally and less in distributed networks, particularly long-range connections. Its is important to keep in mind that these are not absolutes, but rather a relative shift in the balance of how much information is carried in local versus distributed networks.

### Replication

The subjects in the MEG and EEG studies were considered “healthy elderly” by conventional screening. However, a question remains as to whether the changes in complexity reflect truly healthy aging or are indicative of cognitive decline. This was a focus of a follow-up study that examined MSE changes across age in relation to performance on a memory task and lifestyle, particularly physical activity (Heisz, Gould, & McIntosh, 2015).

Participants conducted a directed forgetting task while EEG was measured. Subjects were given a serial list of stimuli and then, based on a cue, are directed to remember all or only part of the list. In addition, to these data, participants completed the Victoria Longitudinal Study Activity Questionnaire and the Montreal Cognitive Assessment battery (MOCA)(Nasreddine, Phillips, & Chertkow, 2012; Nasreddine et al., 2005).

The overall EEG effects in terms of MSE were the same as reported in the original studies (i.e., higher Se at fine scale and less Se at coarse scales for older subjects). There was a trend towards differences in task performance between groups, but was not significant by conventional statistical criteria.

A further assessment of the MSE effects in relation to task performance, physical activity, and MSE scale showed interesting age effects. First, within each group the correlation between fine and coarse scale MSE was calculated showing that in the older group, *greater* Se at fine scale correlated with *less* Se at coarse scales. This correlation was not significant in young subjects.

The scale-dependency was also evident in the correlation with task performance accuracy: higher fine-scale Se and lower coarse scale Se was related to better performance in the older subjects. The dependency in the old subjects is also evident if we plot the MSE curves for the older subjects based on performance split (Figure 9). High performing older subjects showed relatively higher fine- and less coarse-scale Se, relative to younger subjects. Low performing elderly showed less fine-scale Se, compared to high performing elderly, and similar levels of coarse-scale Se as young adults. This scale dependency is also present in relation to cognitive performance on the MOCA, where older adults with worse MOCA scores show less entropy at fine scales and more at coarse scales relative to participants with better MOCA scores.

**Figure 9.**
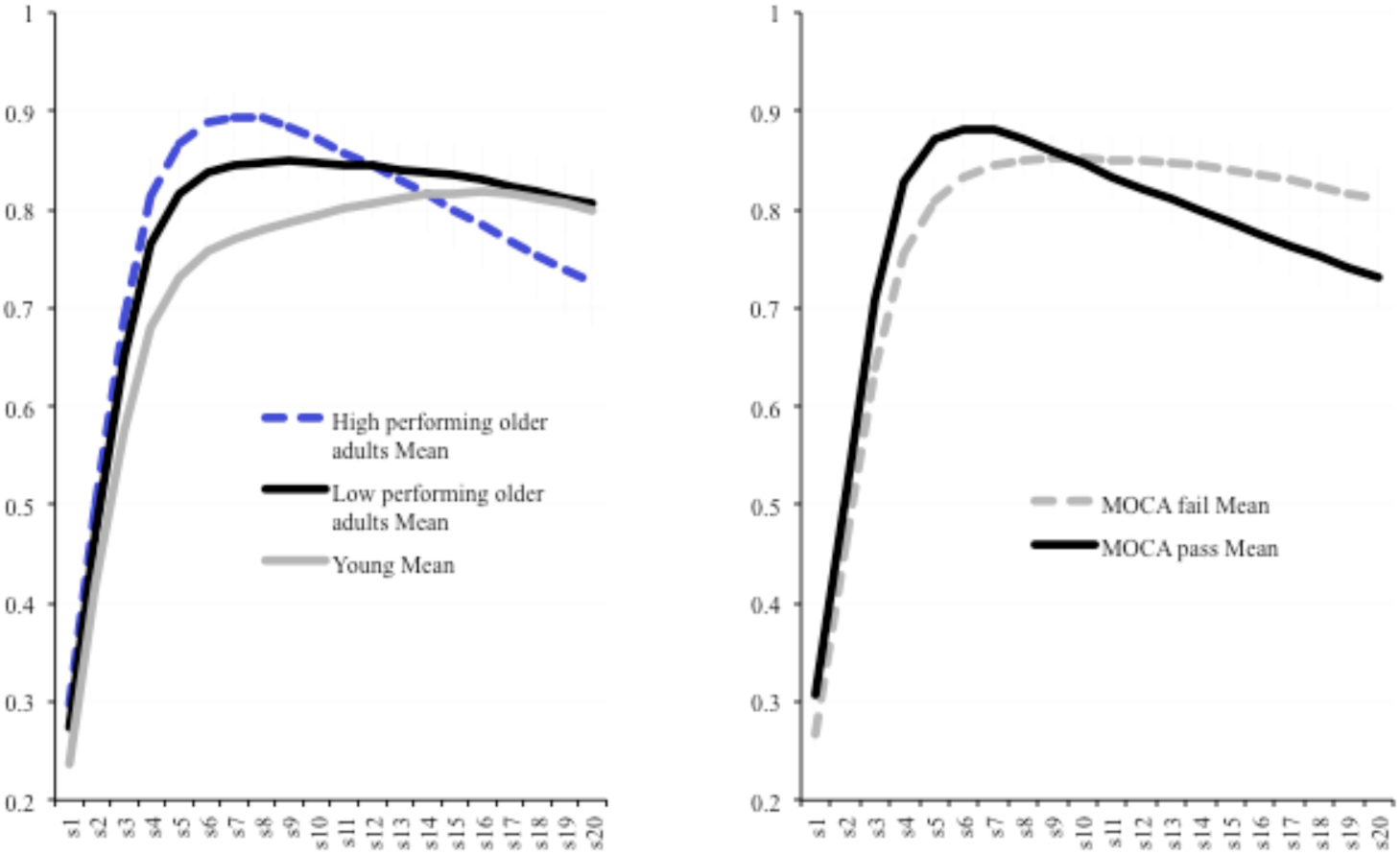
Left panel: MSE curves for young adults (grey) and high- (dashed blue) and low (black) performing old adults. High performing old adults have high fine scale and low coarse scale entropy. Right panel: MSE curves for the same older adults divided base on cognitive performance on the Montreal Cognitive Assessment battery (MoCA). The MSE curve patterns matches that related to performance in the left panel.

Next, within the older subject group, the correlation of MSE scale, task performance and physical activity was examined (with age partialled out). Task performance and physical activity were positively related with Se at fine timescales, whereas only task performance was negatively related to Se at coarse timescales. Physical activity was also positively correlated with great Se at only at fine scale Se.

Finally, a causal modeling analysis was done to ascertain whether the relation of physical activity and task performance could be mediated through the MSE scale changes (Figure 10). This analysis showed both a direct effect of physical activity on task performance, as well as a significant indirect effect going serially from physical activity, to fine scale, Se, coarse scale Se, and then task performance. There was no significant indirect effect if the two MSE scales were considered separately.

**Figure 10.**
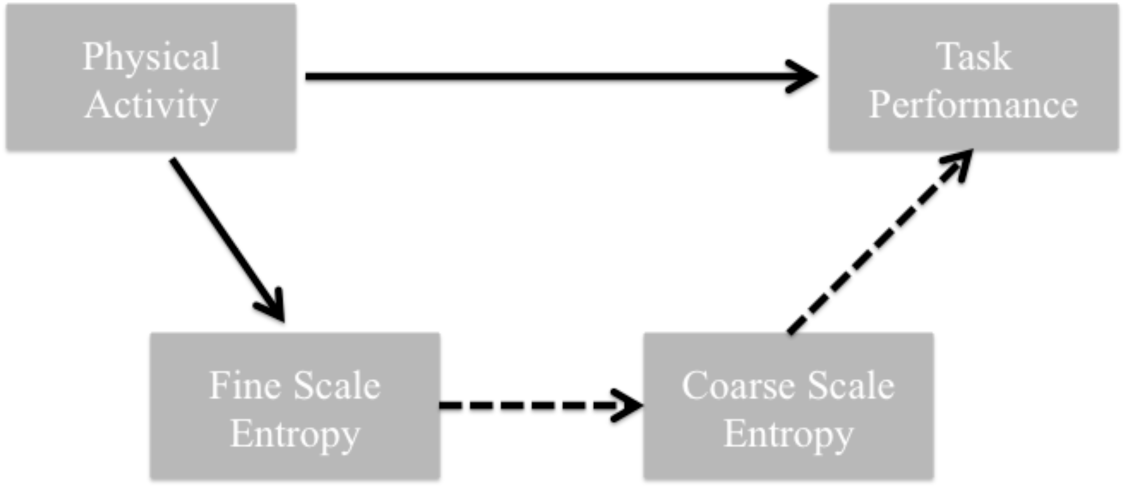
Causal model indicating the impact of physical activity on memory performance and its indirect effect on MSE at fine and coarse scale. Dashed arrows indicate negative effects.

These results have several implications. First, the relation of the MSE scale differences and aging suggest that the relative shift in fine and coarse timescales is, in fact, indicative of healthy aging. The correlation of the fine and coarse scales in the older subjects suggests this balance does vary with healthy aging. Moreover, the relation to physical activity and task performance reinforces the assertion that the fine vs. coarse scale balance can index healthy aging.

Studies that have used fMRI to examine brain signal variance changes in aging have noted spatial variation in these changes (Garrett, Kovacevic, McIntosh, & Grady, 2010). The metric in this study was the standard deviation of the fMRI BOLD timeseries. While many regions show decreased standard deviations with healthy aging, notably medial posterior cortices, there were also some regions that increase in ventral temporal cortices. Task-modulated effects also showed distinct age-related differences, in the face of very similar performance (Garrett, Kovacevic, McIntosh, & Grady, 2013). This reinforces the assertion that age-related changes in complexity are not unidirectional and have a spatiotemporal structure.

#### Clinical extensions

The data from work on healthy aging suggest that scale-dependent changes in brain complexity are expected in the face of relatively good cognitive function (Bertrand et al., 2016). The final study examines how this scale-dependency may act as an index of unhealthy cognitive status. Here we examined persons with Parkinsons Disease (PD), half of which converted to show dementia two years after the initial assessment. There were 62 PD patients (41 men, mean age ± SD: 65.60±8.44 years) and 37 controls (26 men, mean age ± SD = 66.64±8.90 years) included in this study. At follow-up, 44 PD patients were dementia-free (PDnD) and 18 developed dementia (PDD).

Clinical EEG data were taken at the first assessment while the participants were in a resting-state with their eyes-closed but awake. Relative to controls both PD groups showed lower fine-scale entropy. However, the PDD group also showed higher coarse scale Se, relative to both controls and the PDnD group. Keeping in mind these data were collected at least two years prior to the expression of dementia, these finding suggest the higher coarse scale Se may be a predictor of future cognitive decline. The MSE results showed a cohesive link to the healthy aging pattern with respect to the scale-dependent differences, particularlyscales and the scores on the MOCA scale (Figure 11)

**Figure 11.**
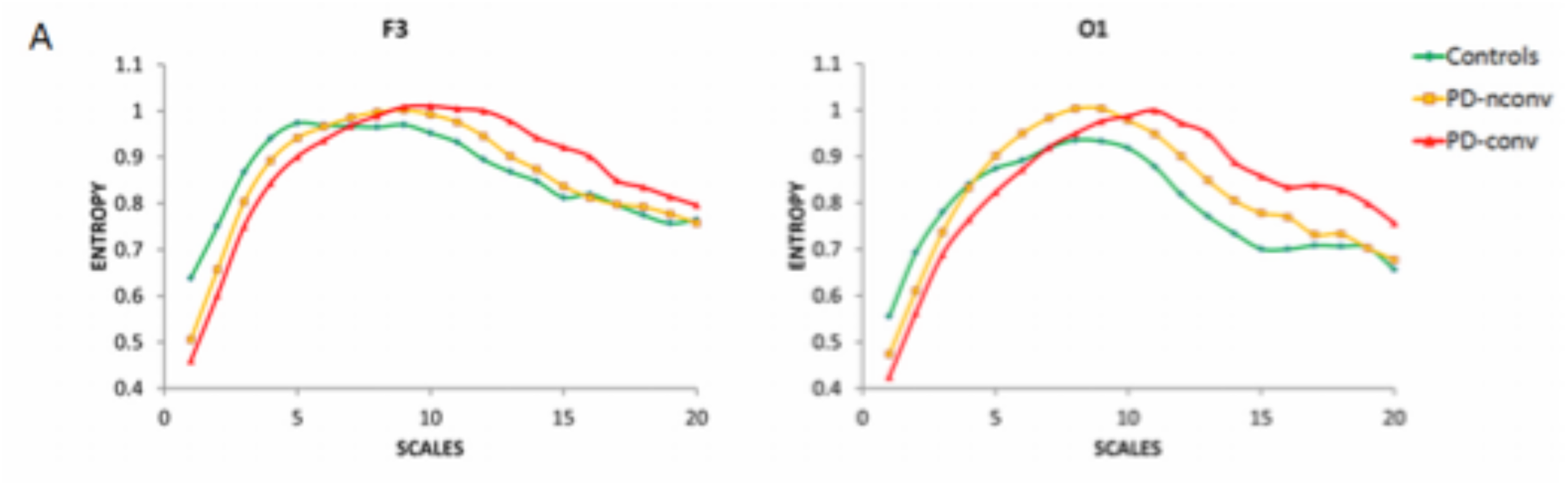
MSE curves from two EEG channels in Parkinson Disease patients (PD) and age-matched controls. PD patients who converted to show dementia (PD-conv) two-years after the initial measure, showed lower fine scale and higher coarse scale entropy relative to non-converters (PD-nconv) and controls.

Other groups have also observed the scale-dependency in dementia. Yang et al (Yang et al., 2013) observed that the scale dependency related with severity of dementia in patients with Alzheimer’s disease, with the morphology of the curve tracking the severity of dementia. Moreover, the fine- vs. coarse-scale balance also related to behavioral symptoms.

It is noteworthy to consider that the specific profile observed in the MSE changes may be related to cognitive impairment as opposed to other disorders. Yang and colleagues have mapped the MSE curves in neuropsychiatric disorders (Hager et al., 2017), noting a mixture of general decreases in complexity and scale-dependent shifts across brain areas. Critically, there is a trend wherein those with psychiatric disorder show regions where there is a move to “randomness”, where fine scale entropy is high followed by a drop that is reminiscent to pure noise processes.

## Conclusions & Future Directions

A general perspective can now be elucidated that summarizes the observations on brain complexity across the lifespan. If we superimpose the space-time structure with the summary MSE curves from the studies that have been reviewed, the observations come as follows:

1. The general morphology of the MSE curve in healthy adults shows increasing entropy with scale, peaking at mid-scales and then showing a slight decline at coarser scales. Adults show the highest information processing (as indexed by entropy) that at a middle scale. This curve can be treated as the standard for subsequent comparisons.
2. Children show overall lower entropy across all scales, and the curve shifts upwards with maturation. The greatest increase is at middle time scales.
3. Healthy old adults show a relative increase of entropy at fine scales and a decline at coarse scales. This maps on to higher local processing and reduced long-range processing.
4. Old adults at risk for cognitive decline do not show the increase at fine scale entropy, and show either similar or higher entropy at coarse scales.

The timescale differences in the curves are suggestive of differing local and global spatial scales over which these effects are expressed (Figure 1). Fine scale entropy would represent local interactions of adjacent cell populations, and the distance of interactions would increase with temporal scale. These mappings are purely speculative, but do represent testable hypotheses that can better relate brain structure and function and the cognitive functions they support.

*Hypothesis 1:* given the relative change in density of the space-time structure of the connectivity matrix, and the morphology of the MSE curves, the highest information exchange in the brain should occur at intermediate spatiotemporal scales. This could be assessed through estimates of mutual information, or comparable, as a function of connection distance with the expectation that information would be highest for the middle distance connections

*Hypothesis 2:* the relative shift to faster timescale (local) processing indicates healthy brain function. While this is a simple restatement of the empirical findings, if this shift is, in fact, an accurate index of healthy aging, it could impact how we consider the neural support for cognition.

We have observed in healthy adults that episodic memory processes tend to emphasize local processes more so that semantic memory (Heisz, Vakorin, Ross, Levine, & McIntosh, 2014). If this relation is maintained in healthy aging, then it suggests that the nature of cognitive processes associated with these memory functions may show a qualitative change, despite being under the guise of “healthy aging”.

For this second hypothesis, there would be tremendous benefit in a longitudinal assessment of the observed scale-dependent entropy shifts. The MSE curves presented here reflect group averages and snap-shots in time, so it is not possible to ascertain how much of what we see reflects a current “state” of ones brain, or a “trait” that may be a stable index of information processing capacity.

The space-time changes that we report also have obvious clinical utility as an early marker of risk for cognitive decline. Again, longitudinal testing would determine whether the space-time pattern is a trait or shows changes across time.

I have used the notion of complex adaptive systems to review work, primarily from my own lab, that characterizes brain complexity changes in aging. These changes can be conceptualized as a reflection of changing information processing capacity. An important aspect of this change is that it reflect changes in space and time, hence “which” areas/networks and “when” for a given cognitive process may indeed change as we age. There are potential implications for the nature of the link between brain and behavior if we assume that the elemental processes that are linked in coordinating behavior show the same features of complexity as the brain (Kelso, 1995; Perdikis, Huys, & Jirsa, 2011; Pillai & Jirsa, 2017). This could indeed bring a different perspective on the fundamental cognitive processes where “cognitive deficits” in aging is not simply a decline of the system that we had as young adults, but rather a reconfiguration or adaptation of the system. Thus, in healthy aging, at least the changes are a natural evolution of the brain rather that a non-optimal state. In this framework, the perspective on disease and degeneration also changes in that the overall behavior that emerges is an adaptation of the system. The accompanying deficits are thus as much a reflection of the damaged or missing elements, as they are the expression of a new repertoire of the system following adaptation.

